# AcuM and AcuK: the global regulators controlling multiple cellular metabolisms in a dimorphic fungus *Talaromyces marneffei*

**DOI:** 10.1101/2024.04.10.588979

**Authors:** Tanaporn Wangsanut, Artid Amsri, Thitisuda Kalawil, Panwarit Sukantamala, Juthatip Jeenkeawpieam, Alex Andrianopoulos, Monsicha Pongpom

## Abstract

Talaromycosis is a fungal infection caused by an opportunistic dimorphic fungus *Talaromyces marneffei*. During infection, *T. marneffei* resides inside phagosomes of human host macrophages where the fungus encounters nutrient scarcities and host-derived oxidative stressors. Previously, we showed that the deletion of *acuK,* a gene encoding Zn(2)Cys(6) transcription factor, caused a decreased ability for *T. marneffei* to defend against macrophages, as well as a growth impairment in *T. marneffei* on both low iron-containing medium and gluconeogenic substrate-containing medium. In this study, a paralogous gene *acuM* was deleted and characterized. The Δ*acuM* mutant showed similar defects with the Δ*acuK* mutant, suggesting their common role in gluconeogenesis and iron homeostasis. Unlike the pathogenic mold *Aspergillus fumigatus*, the Δ*acuK* and Δ*acuM* mutants unexpectedly exhibited normal siderophore production and did not show lower expression levels of genes involved in iron uptake and siderophore synthesis. To identify additional target genes of AcuK and AcuM, RNA-sequencing analysis was performed in the Δ*acuK* and Δ*acuM* strains growing in a synthetic dextrose medium with 1% glucose at 25 °C for 36 hours. Downregulated genes in both mutants participated in iron-consuming processes, especially in mitochondrial metabolism and anti-oxidative stress. Importantly, the Δ*acuM* mutant was sensitive to the oxidative stressors menadione and hydrogen peroxide while the Δ*acuK* mutant was sensitive to only hydrogen peroxide. The yeast form of both mutants demonstrated a more severe defect in antioxidant properties than the mold form. Moreover, ribosomal and ribosomal biogenesis genes were expressed at significantly lower levels in both mutants, suggesting that AcuK and AcuM could affect the protein translation process in *T. marneffei*. Our study highlighted the role of AcuK and AcuM as global regulators that control multiple cellular adaptations under various harsh environmental conditions during host infection. These transcription factors could be potentially exploited as therapeutic targets for the treatment of this neglected infectious disease.

**AUTHOR SUMMARY:** *Talaromyces marneffei* invades host macrophages to establish infection. Major stressors inside the macrophage compartments are nutrient deprivation and oxidative substances. Here, we demonstrated that AcuK and AcuM transcription factors are necessary for *T. marneffei* to grow under iron and glucose limitation, and to survive oxidative stress and macrophage killing. AcuK and AcuM regulate non-glucose carbon utilization via the transcriptional control of gluconeogenic genes. For iron homeostasis, the two proteins regulate the expression of genes involved in iron-utilization pathways. Lastly, the AcuK and AcuM play a role in oxidative stress response likely by regulating the expression of genes encoding antioxidant enzymes and alternative respiration enzymes. Thus, AcuK and AcuM control multiple cellular adaptations that allow *T. marneffei* to cope with major stressors occurring during macrophage infection. Since AcuK and AcuM are critical for cellular metabolism and macrophage engulfment, this new information could lead to a better understanding of host-pathogen interaction and could be ultimately developed into fungal-specific diagnostic tools and therapeutic agents.

## INTRODUCTION

Talaromycosis is caused by the opportunistic fungal pathogen *Talaromyces marneffei,* which mainly affects immunocompromised individuals (1). It is commonly widespread in patients with HIV/AIDS as well as patients with defects in cellular immunity, especially in CD4 T-cell activity (2, 3). *T. marneffei* infections are endemic in Southeast Asia and South China with over 17,300 cases reported annually (4). Among infected individuals, up to one-third are associated with death (5). Although talaromycosis is strongly associated with high morbidity and mortality, and *T. marneffei* is listed as one of the fungal priority pathogens by the World Health Organization (WHO), talaromycosis still has not received sufficient attention and investment from regional and global funders, researchers, clinicians, industries, and policy makers (4, 6). As a result, the control and prevention of this neglected infectious disease remains difficult.

*T. marneffei* is a thermal dimorphic fungus that grows as a mold at environmental temperatures (25°C) and undergoes morphological switching to fission yeast at human body temperature (37°C) (7). Conidia, the infectious asexual spores from the environment, can be inhaled into a patient’s lungs and subsequently engulfed by alveolar macrophages, where the conidia switch to the pathogenic yeast cells and cause infection (8, 9). The fungus resides within phagosomes of macrophages, and therefore *T. marneffei* is a facultative intracellular pathogen. However, the host macrophage compartments present several harsh environmental conditions such as nutrient scarcities and oxidative stressors to *T. marneffei* (10). Thus, the pathogenicity of *T. marneffei* is dependent on its ability to retrieve sufficient nutrients, survive host-derived stress, and replicate inside the macrophage cells.

Gluconeogenesis and iron metabolism are crucial for the establishment of *T. marneffei* infection. The macrophage phagosome, where the microorganism is engulfed and resides, typically contains gluconeogenic substrates. Presumably, the capacity to metabolize these substrates for use in gluconeogenesis is crucial for energy production and intracellular growth of *T. marneffei* (10, 11). Additionally, the human host limits iron availability via nutritional immunity mechanisms. To cope with iron scarcity, *T. marneffei* has evolved high-affinity iron uptake systems, including reductive iron assimilation (RIA) and siderophore-assisted iron uptake (12). Following iron acquisition, iron is utilized by several important cellular pathways, generally as the cofactor in the form of Fe-S clusters, or as the center of heme groups. Iron-dependent processes include the tricarboxylic acid (TCA) cycle, the electron transport chain, lipid and sterol metabolism, biotin and lipoic acid cofactor synthesis, amino acid biosynthesis, DNA replication and repair, chromatic remodeling, and protein translation (13). As such, *T. marneffei* requires sufficient amounts of glucose and iron to ensure survival inside host cells; therefore, gluconeogenesis and iron metabolic processes are required for pathogenesis (12, 14).

Microorganisms generally have evolved a set of regulatory proteins that can modulate their metabolic adaptation (13, 15, 16). Transcriptional control of carbon metabolic genes plays an important role in maintaining homeostasis during the switch from one carbon substrate to another (16). In the mold model *Aspergillus nidulans*, AcuK and AcuM are homologous transcription factors that regulate the expression of genes specific for gluconeogenesis and the TCA cycle (16, 17). The *acuK* and *acuM* genes were originally identified by the selection for *A. nidulans* mutants defective in acetate utilization (18). Subsequently, these two factors were shown to directly bind the promoters of gluconeogenic genes *acuF* (phosphoenolpyruvate carboxykinase, PEPCK) and *acuG* (fructose-1,6-bis phosphatase, FBP) genes to activate their expression (16, 19). AcuK and AcuM proteins contain the Zn(2) Cys(6) binuclear cluster DNA-binding domain, and function *in vivo* in the form of heterodimers (17). These transcription factors are fungal-specific and found only in Ascomycete filamentous fungi. Surprisingly, studies in the pathogenic mold *Aspergillus fumigatus* have revealed an additional role of AcuK and AcuM in iron acquisition and virulence (20, 21). SreA/HapX are two well-known transcription factors that control the expression of genes in the RIA (*fre1* and *fre2*) and siderophore-assisted uptake (*sidA*, *sidD*, *mirB*) systems (22). To efficiently respond to different levels of iron, SreA and HapX can regulate expression levels of each other in a negative feedback loop manner (23). In fact, in *A. fumigatus* AcuK and AcuM control the iron assimilation process through the inhibition of the SreA transcription factor (24). Besides the studies in *A. fumigatus,* the target genes under the control of AcuK and AcuM have never been fully investigated in other pathogenic fungi.

The *T. marneffei* genome contains orthologous genes that encode for AcuK and AcuM. Previously, we characterized the function of the *acuK* homologue in *T. marneffei* (25). The *acuK* deletion mutant showed growth defects under low iron conditions and was unable to utilize and grow on gluconeogenic carbon sources (25). Also, the Δ*acuK* mutant exhibited increased killing by THP-1 human macrophage cells. In this study, we investigated the role of AcuM in iron and carbon metabolism by generating the Δ*acuM* mutant and characterizing its phenotypes. As seen in the Δ*acuK* mutant, the Δ*acuM* mutant showed strong growth reduction in media containing low iron levels and gluconeogenic carbon sources. To identify the global AcuK and AcuM target genes, transcriptome profiles were analyzed using an RNA-sequencing approach. Unexpectedly, we discovered that AcuK and AcuM did not directly control the genes in RIA and siderophore biosynthesis as found in *A. fumigatus*, but rather they affected the expression of genes in iron-consuming pathways. Specifically, the AcuM-dependent and AcuK-dependent genes encoded iron-containing proteins that function in mitochondrial carbon metabolism, protein synthesis, and oxidative stress response. The Δ*acuM* mutant showed higher sensitivity to menadione and hydrogen peroxide than the Δ*acuK* mutant. Lastly, deletion of the *acuM* gene attenuated *T. marneffei* virulence in a macrophage infection model. This study demonstrated that AcuK and AcuM are important regulators required for adaptation to and defense against the harsh environmental conditions presented by host macrophages during infection.

## MATERIALS AND METHODS

### 1. Fungal Strains and Culture Conditions

*Talaromyces marneffei* ATCC18224 (FRR2161) is used as the wild type strain in this study. A G809 strain (Δl*igD pyrG^+^ niaD^-^*) was used as the *acuM*+ strain in the macrophage infection assay. The mutant strains, Δ*acuK* (Δ*acuK Anpyr*G+) and Δ*acuM* (Δ*acuM AnpyrG+*) were generated from a background of the uracil auxotrophic G816 (Δl*igD pyrG^-^ niaD^-^*) strain (26). *T. marneffei* G816 strain was cultured on *Aspergillus* minimal medium (ANM; 1% glucose, mixture of trace elements and 10 mM (NH_4_)_2_SO_4_) and supplemented with 5 mM uracil. *T. marneffei* wild type, G809 (*acuM*^+^), Δ*acuM*, and Δ*acuK* strains were cultured on the ANM without uracil. Conidia were harvested from a 10-day-old culture on a solidified agar plate by scraping the colony surface and resuspending in a sterile normal saline-tween solution (0.1% v/v Tween 40, 0.85% w/v NaCl). The suspension was thoroughly vortexed and filtered through sterile glass wool to remove the mycelia. The conidial concentration was enumerated by counting in a hemacytometer.

### 2. Construction of *acuK* and *acuM* deletion strains

The *acuK* deletion strain (Δ*acuK*) was constructed previously (25). The *acuM* deletion strain (Δ*acuM*) was generated in this study. The *acuM* open reading frame and flanking regions were amplified from the genomic DNA of the FRR2161 strain with primers pair 5’-AcuM-F-*Apa*I (5′-ATGGGCCCAGAGATTGCCGCGGTCCTAC-3′) and 3’-AcuM-R-*Sac*I (5′-GCGGAGCTC TCACCATCTCCACAGCTGTGTG-3′). The 5.3-kb product was cloned into pGEM-T Easy (Promega), generating an *acuM* cloning plasmid, pGEMT-*acuM*. To generate an *acuM* deletion construct, a pGEM-T plasmid backbone including 5’ and 3’ flanking regions without *acuM* coding region was amplified by inverse PCR with inv_3_*Eco*RV (5’-CGCAGCAGATATCATGTCCC-3’) and inv_5_SmaI (5’-ATATCCCGGGGGCGTTCTTGAGCACAAGT-3’). The 4.8-kb product was ligated with an *Aspergillus nidulans pyrG* (*AnpyrG*) selectable marker, which was digested with *Eco*RV and *Sma*I from plasmid pAB4626, producing the *acuM* deletion plasmid, pDel_*acuM*. A deletion cassette composed of *AnpyrG* flanks with *acuM* 5’ and 3’ regions was amplified from the pDel_*acuM* deletion plasmid. Transformation of the deletion cassette into the G816 protoplasts was performed by using a PEG/CaCl_2_ method as described previously (26).

### 3. Determination of growth in iron-deplete and iron-replete conditions

To study the effect of iron availability on the growth of *T. marneffei*, five microliters of conidial suspension containing 10^5^ conidia/ml were spotted on the surface of ANM medium with different iron concentrations: normal (ANM, 7 µM ferrous sulfate), iron deplete (ANM, 100 µM phenanthroline) and iron replete (ANM, 100 µM phenanthroline and 1 mM FeCl_3_). The colony was observed after incubation for 7 days at either 25 °C or 37 °C. Measurement of siderophore production was performed as previously described (27).

### 4. Determination of growth on medium containing gluconeogenic substrates

A carbon-free agar (CF) medium (mixture of trace elements and 10 mM (NH_4_)_2_SO_4_) was prepared. The gluconeogenic carbon substrates were added as follows: 50 mM proline, 50 mM acetate, or 0.5% ethanol. As a control, 1% glucose was added to the CF medium. The conidial suspension was harvested from the colonies of *T. marneffei* wild type, Δ*acuK*, and Δ*acuM* strains. Five μl containing 10^5^ conidia of each strain were spotted on the surface of the medium and incubated at either 25 °C or 37 °C. Colonies were observed after incubation for up to 14 days.

### 5. Determination of growth on medium containing oxidative stressors

Oxidative stress susceptibility tests used an agar drop dilution assay. Five microliters containing conidia of *T. marneffei* ATCC18224, Δ*acuK*, or Δ*acuM* strains were spotted on the surface of ANM medium containing oxidative stressors, hydrogen peroxide (1, 2,3 and 5 mM) and menadione (5 and 10 μM). Due to the differences in growth rate, the inoculum conidial numbers were 10^2^ - 10^4^ for the mycelium growth assay and 10^5^ - 10^7^ for the yeast growth assay. The plates were incubated at either 25 °C or 37 °C for up to 14 days. Colonies were observed and photographed.

### 6. Quantitative RT-PCR (qRT-PCR)

*T. marneffei* ATCC18224, Δ*acuK,* and Δ*acuM* strains were prepared to 10^8^ conidia/ml in 100 ml ANM medium and were pre-cultivated at 25 °C for 16 hours to generate germinating conidia. The fungal cells were washed three times with sterile PBS before being transferred into a new medium for different test conditions. For detection of the gluconeogenic genes (list of gene names and primers used, Table S1), the pre-cultivated fungal cells were transferred to the CF medium containing the following gluconeogenic carbon sources; 50 mM proline, 50 mM acetate, and 0.5% ethanol. To investigate the expression of genes in the iron assimilation pathways (Table S1), the pre-culture was transferred to the iron-free ANM broth supplemented with different iron concentrations (0 – 1000 mM FeCl_3_). The cultures were incubated at either 25 °C for 24 hours or 37 °C for 48 hours before harvesting the fungal cells by centrifugation, and the RNA was extracted from the cells.

Two micrograms of the isolated total RNA were converted to cDNA using ReverTra Ace qPCR RT Master Mix (TOYOBO Inc., Osaka, Japan). The qRT-PCR was performed using the SYBR green qPCR mix (Thunderbird SYBR green Chemistry, Toyobo, Osaka, Japan) and the intensity of the fluorescent signal was detected using a 7500fast real-time PCR system (Applied Biosystems, Foster, CA, USA). The primers of selected genes are shown in Table S1. The expression level of each gene was tested in the following conditions: one cycle of 95°C, 60 sec; followed by 40 cycles of 95°C, 60 sec and 60°C, 60 sec. Comparative quantification (fold change) levels were calculated by the comparative cycle threshold (2^-ΔΔCt^) method, and the actin transcript was used as an internal control.

### 7. RNA sequencing

To prepare samples for the RNA-seq experiment, conidia of *T. marneffei* ATCC18224, Δ*acuK* and Δ*acuM* were inoculated at a final concentration of 10^8^ cell/mL in 250 mL synthetic dextrose (SD) broth (1% w/v glucose, 0.17% w/v yeast nitrogen base without amino acids, 10 mM (NH_4_)_2_SO_4_,) and incubated at 25°C for 36 hours for the generation of mycelium mass. The cultures were harvested by centrifugation at 4,500 rpm for 10 min. Approximately 2 g wet weight was homogenized in a TRIzol reagent (Invitrogen, Life Technologies, Carlsbad, CA, USA) by using a bead beater (Biospec, Bartlesville, OK, USA). Total RNA was isolated following the manufacturer’s protocol. The RNA was digested with 1 U/µL DNase I to eliminate the residual DNA and purified by isopropanol precipitation. RNA concentrations were determined with a spectrophotometer (Nanodrop 2000: Thermo Scientific, Waltham, MA, USA). RNA integrity and purity were tested by agarose gel electrophoresis. Library preparation and RNA sequencing were performed by a service from Novogene Co., Ltd. (Hong Kong). Messenger RNA was purified from total RNA using poly-T oligo-attached magnetic beads. After fragmentation, the first strand cDNA was synthesized using random hexamer primers, followed by the second strand cDNA synthesis using either dUTP for directional library or dTTP for non-directional library. Libraries underwent end repair, A-tailing, adapter ligation, size selection, amplification, and purification. Libraries were checked with a Qubit Fluorometer and real-time PCR for quantification and bioanalyzer for size distribution detection. Quantified libraries were pooled and sequenced on Illumina NovaSeq6000 machine to generate the paired-end reads.

Sequencing reads were annotated and aligned to the *T. marneffei* ATCC18224 reference genomes by using Hisat2 v2.0.5. StringTie (v1.3.3b) was used to assemble the mapped reads and predict novel transcripts in a reference-based approach (28). To quantify gene expression level, the feature-Counts v1.5.0-p3 was used to count the read numbers mapped to each gene and then FPKM (Fragments Per Kilobase of transcript per Million mapped reads) for each gene was calculated based on the length of the gene and reads count mapped to the gene. To analyze differential gene expression in each sequenced library, the read counts were adjusted using the edgeR package through one scaling normalized factor. Differential expression analysis of two conditions was performed using the edgeR R package (3.22.5). The P values were adjusted using the Benjamini & Hochberg method. Corrected P-value of 0.05 and an absolute fold change of 2 were set as a threshold for significant differential expression. Gene Ontology (GO) enrichment analysis and metabolic pathway enrichment analysis of differentially expressed genes were performed by the cluster Profiler R package. GO terms with corrected P-value less than 0.05 were considered significantly enriched by differentially expressed genes. The Cluster Profiler R package was used to test the statistical enrichment of differential expression genes in KEGG pathway. The RNA-seq data were deposited and available at Gene Expression Omnibus (GEO) accession GSE248925.

### 8. Macrophage infection for phagocytosis and killing assay

The THP-1 human monocytic cell line was used for a macrophage infection model, with cells maintained in RPMI 1640 medium (Life Technologies, Grand Island, USA) with 10% FBS (v/v) at 37 °C, 5% CO_2_. THP-1 cells (1 × 10^6^) were inoculated into each per well of a 6-well microtiter plate containing one sterile coverslip for phagocytosis assay and a 12-well plate for killing assay. The cell was differentiated with 32 µM phorbol 12-myristate 13-acetate (PMA) (Sigma-Aldrich) for 24 h at 37 °C and 5% CO_2_. Then, 1 × 10^6^ *T. marneffei* conidia of G809 (*acuM*+) and Δ*acuM* were added to the macrophages and infected for 2 h. For the phagocytosis assay, macrophages were fixed with 4% paraformaldehyde and stained with 1 mg/mL calcofluor white to observe the fungal cell walls. Mounted coverslips were checked using differential interference contrast (DIC) and epifluorescence optics for cell wall staining and viewed on a Reichart Jung Polyvar II microscope. Pictures were captured using a SPOT CCD camera (Diagnostic Instruments Inc., Sterling Heights, MI, USA). The number of phagocytosed cells was recorded in a population of approximately 100 macrophages in three independent experiments. The phagocytic index (the number of phagocytosed conidia per macrophage) was determined by dividing the total number of intracellular conidia by the number of macrophage cells containing conidia. For the killing assay at 2 h after inoculation, the macrophage cells were harvested and lysed with 0.25% TritonX-100 (Sigma-Aldrich). The macrophage killing at 24 h was also determined. After 2 h of infection, the macrophage cells were washed three times with PBS, and the infection was continued for an additional 24 hours before cell lysis. The recovered fungal cells were plated on a synthetic dextrose agar (SD; 0.17% w/v yeast nitrogen base without amino acids, 2% w/v glucose, 10 mM (NH_4_)_2_SO_4_, 2% agar). Colony forming units (CFU) was determined after growth at 37 °C for 7–10 days. Data were expressed as the mean and standard error of the mean. from triplicate experiments.

### 9. DNA binding motif analysis

Prediction of the target genes of AcuK and AcuM in *T. marneffei* was performed by bioinformatic analysis against *T. marneffei* genome. One thousand base pairs upstream regions of the genes in *T. marneffei* ATCC18224 genome were scanned for putative AcuK and AcuM binding motif, CCGNNNNNNNCCG, using the web-based tools available at https://meme-suite.org/meme/tools/fimo (29). One thousand matches with the significant p-values less than 0.0001were subjected to functional enrichment analysis, using DAVID available at https://david.ncifcrf.gov/home.jsp (30). Otherwise, a manual survey for the CCGNNNNNNNCCG motif was carried out in Microsoft Word within the 1,000 bp region upstream of the start codon, ATG for specific target genes.

### 10. Statistical Analysis

Statistical analysis of the data depending on the experiment and included a Student’s t-test and Tukey’s multiple comparison test or unpaired t-test and Welch’s correction with a significant value of p < 0.05. All statistical analysis was performed by using SPSS v. 16.0 and Prism software (GraphPad, version 7.0).

## RESULTS

### AcuK and AcuM are required for normal growth of *T. marneffei* in medium containing gluconeogenic carbons

In *A. nidulans*, AcuK and AcuM form a heterodimer to regulate the expression of genes in the gluconeogenesis pathway (16). Mutations in the *acuK* or *acuM* genes in *A. nidulans* and *A. fumigatus* halt growth under gluconeogenic carbon sources (17, 20, 21). We previously showed that *acuK* controls the growth of *T. marneffei* on acetate, ethanol, and proline gluconeogenic substrates (25). To investigate if *acuM* was also required for utilization of non-glucose carbon sources in *T. marneffei,* we generated an *acuM* deletion mutant and examined its phenotypic effects, specifically the growth on solid medium containing acetate, ethanol, or proline as the sole carbon source. The Δ*acuK* mutant was included in the experimental design to allow for comparison of the phenotypes between each mutant. Deletion of the *acuM* gene did not cause morphology defects when glucose was sufficient (Fig 1A and E). However, like Δ*acuK*, the Δ*acuM* mutant showed severe growth defects on media containing each of the tested gluconeogenic substrates (Fig 1). Notably, the severity of the growth defect was more prominent during the yeast phase (Fig 1E-H) compared to the hyphal phase (Fig 1A-D). This indicates firstly, that both *acuK* and *acuM* genes are important for growth of *T. marneffei* when glucose is unavailable as a carbon source, especially in the pathogenic yeast form. Secondly, this indicates that the two genes are not redundant.

**Figure 1.**
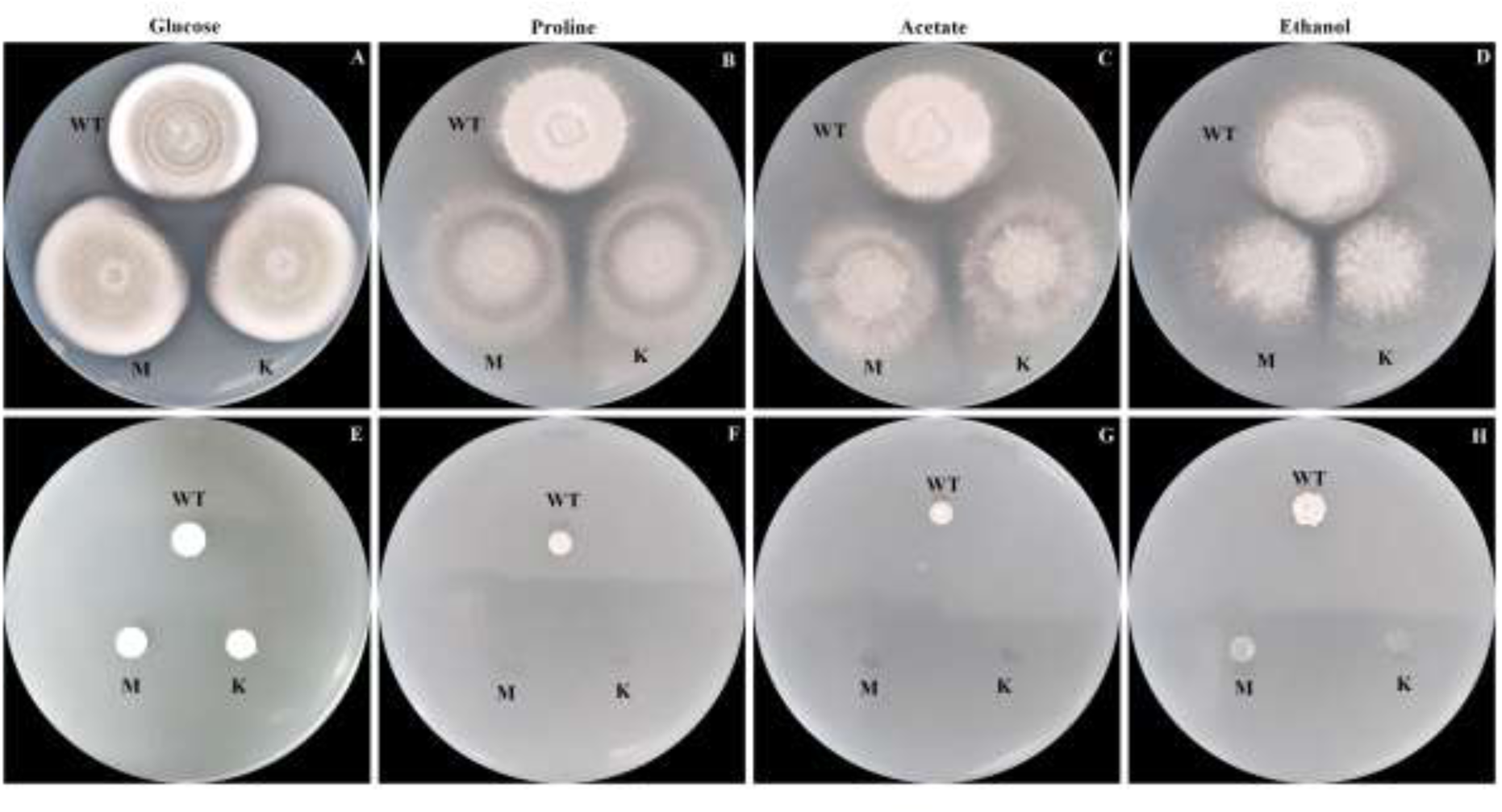
Growth of *T. marneffei* strains were determined on gluconeogenic substrates-containing medium. Conidia suspensions of the *acuK*Δ, *acuM*Δ, and wild type (ATCC18224) strains were prepared at a concentration of 10^8^ conidia/mL. Five microliters of each conidial suspension were spotted onto the surface of the medium supplement with glucose (A and E), proline (B and F), acetate (C and G) and ethanol (D and H). At 25 °C, the mold colony of wild-type, Δ*acuK* and Δ*acuM* strains were observed at 12 days (A – D). At 37 °C, the yeast colony was observed at 12 days (E – H). M = *acuM* deletion mutant; K = *acuK* deletion mutant. Experiments were performed in three biological replicates.

### AcuK and AcuM are required for optimal expression of genes in gluconeogenic carbon utilization

Target genes of *T. marneffei* in the gluconeogenesis pathway that were possibly controlled by AcuK or AcuM were selected by using homologs to the known AcuK and AcuM targets identified in *A. nidulans* and *A. fumigatus*. These were *fbpA* (fructose-1,6-bis phosphatase, a key enzyme in gluconeogenesis), *prnD* (proline dehydrogenase, a key enzyme in proline utilization), *alcB* (alcohol dehydrogenase, a key enzyme in ethanol utilization) and *facB* (a transcription factor for acetate utilization). Presence of the putative DNA binding motif of AcuK and AcuM (CCGN7CCG) homologs from *A. nidulans* (17) was scanned in the upstream region of selected genes and results were listed in Table S2.

*T. marneffei* conidia (10^8^ conidia/ml) were pre-cultured in a liquid ANM medium at 25 °C for 16 hours before being transfered to a carbon-free medium supplemented with 50 mM proline, 50 mM acetate, or 0.5% ethanol to serve as the alternative carbon source. The cultures were incubated for an additional 24 hours at 25 °C or 48 hours at 37 °C. The RNA was extracted from the harvested cells and the transcription levels were assessed by qRT-PCR. Notably, the mutants grew poorly under the non-glucose conditions at 37 °C and we were unable to obtain sufficient RNA samples, so the expression levels were determined only in the mycelium form. In the wild type, the *fbpA* transcript was upregulated the most when the strain was grown in ethanol (Fig 2A). However, the *acuM* mutant did not upregulate *fbpA* gene expression as efficiently as the wild type (40-fold in wild type vs 18-fold in *acuM* mutant). The *acuK* mutant did not show the *fbpA* gene regulation defect when compared to the wild type. Thus, AcuM, but not AcuK, activates the *fbpA* gene and allows cells to utilize ethanol as a sole carbon source. Notably, the wild type strain did not regulate the *fbpA* gene in response to acetate and proline as carbon sources. Nonetheless, the *fbpA* gene was expressed at higher levels in both *acuK* and *acuM* mutants. This result suggests that AcuK and AcuM could be repressors of the *fbpA* gene or AcuK and AcuM could indirectly regulate the *fbpA* gene when acetate and proline are available as sole carbon substrates.

**Figure 2.**
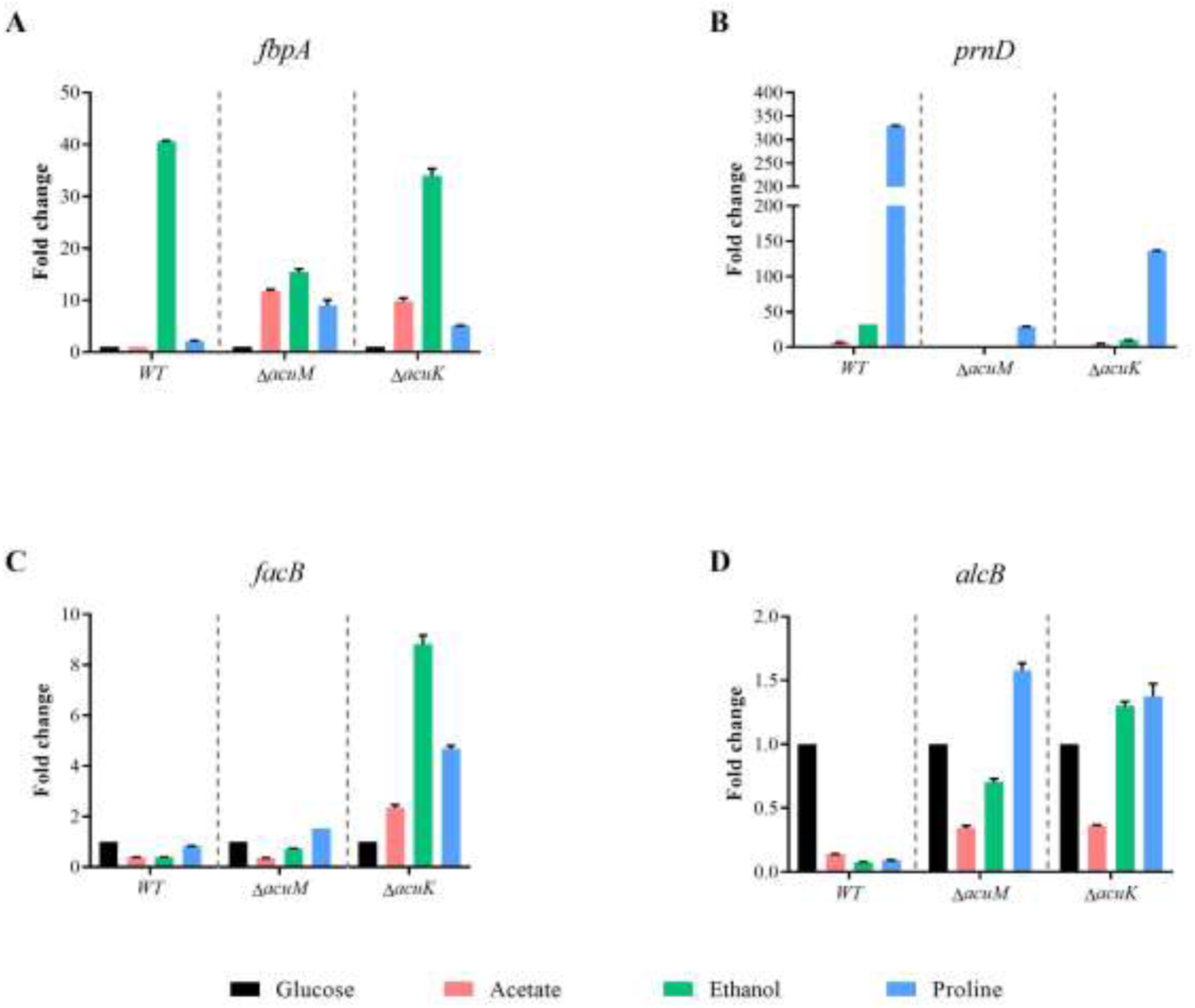
Expression levels of genes involved in carbon metabolism were analyzed in the *acuK* and *acuM* deletion mutants. The 10^8^ conidia/ml of *T. marneffei* wild type, Δ*acuK,* and Δ*acuM* strains were precultured in the minimal medium containing glucose for 16 hours before transferring into acetate, ethanol and proline gluconeogenic carbon source medium. After incubation at 25 °C for 24 hours, the fungal cells were collected and subjected for RNA extraction and cDNA synthesis. The qRT-PCR was used to determine the fold-change of gene transcripts (A) *fbp* (fructose-1,6-bis phosphatase), (B) *prnD* (proline dehydrogenase), (C) *facB (*key regulator in acetate utilization), and (D) *alcB* (alcohol dehydrogenase). Gene expression levels were calculated by the 2^-ΔΔCt^ method using actin as a reference gene.

When proline was used as a sole carbon source, the *prnD* gene was upregulated 320-fold in the wild type (Fig 2B). However, the *prnD* gene was expressed at lower levels in both mutants, showing a 10-fold decrease in the *acuM* mutant and 2-fold decrease in the *acuK* mutant, compared to the wild type (Fig 2B). Thus, AcuM and AcuK could activate the *prnD* gene to allow the cells to metabolize proline. AcuM seemed to play a more prominent role than AcuK as the *acuM* mutant showed more severe gene regulatory defects than the *acuK* mutant.

While *facB* gene expression levels were not transcriptionally induced in response to acetate, ethanol, or proline as alternative carbon sources, the *acuK* mutant expressed the *facB* gene at higher levels, exhibiting a 3-fold, 9-fold, and 5-fold increase in response to acetate, ethanol, and proline, respectively (Fig 2C). The *acuM* mutant showed a similar gene expression pattern as the wild type, indicating that AcuM was not involved with transcriptional regulation of the *facB* gene. This result implies that AcuK could be a repressor of the *facB* gene or AcuK indirectly regulates the *facB* gene under tested growth conditions.

In the wild type, the *alcB* gene expression was transcriptionally repressed when acetate, ethanol or proline was added as the sole carbon source (Fig 2D). However, the *alcB* gene was upregulated in both *acuM* and *acuK* mutants, especially when the cells were transferred into medium containing ethanol or proline. This result suggests AcuM and AcuK could repress the expression of the *alcB* gene or AcuM and AcuK could indirectly regulate the *alcB* gene under tested growth conditions.

Altogether, our data suggests that AcuK and AcuM regulate the expression of genes required for gluconeogenic carbon utilization to varying degrees. Both AcuM and AcuK are involved in the regulation of *alcB* gene expression when gluconeogenic substrates are available as sole carbon sources. AcuM played a prominent role in regulating the *fbpA* and *prnD* gene expression while AcuK affected the expression of the *facB* gene. In addition, AcuK and AcuM could potentially function as either activators or repressors to control the target genes in gluconeogenesis pathways.

### AcuK and AcuM are required for growth in low iron conditions

AcuK and AcuM have evolved the additional function of controlling high-affinity iron acquisition in *A. fumigatus* (20, 21). In *T. marneffei,* AcuK is necessary for growth under iron limitation (25). To investigate whether AcuM is also involved in *T. marneffei* iron regulation, we examined the growth of the wild-type and Δ*acuM* strains under various iron concentrations. Strains were grown on ANM solid medium, defined as normal (containing 7 µM of iron), iron-deplete (ANM with 100 µM phenanthroline) and iron-replete (ANM with 100 µM phenanthroline and 1 mM FeCl_3_) conditions (Fig 3). Experiments were performed during both hyphal and yeast phase growth and the Δ*acuK* mutant was included to permit phenotypic comparisons between each strain. At 25°C, the hyphal colony morphology of the Δ*acuM* and Δ*acuK* mutants was not different from the wild type strain under both normal and iron-replete conditions (Fig 3, A and C). The mutants showed normal morphology with grayish green coloration and a velvety to powdery colony surface. The growth of all strains was higher in iron replete conditions than in normal conditions, indicating that iron enrichment enhanced growth of the fungus (Fig 3C). However, the Δ*acuK* and Δ*acuM* strains were unable to grow when iron was depleted in both hyphal and yeast phases (Fig 3B). These results indicate that AcuK and AcuM are indispensable for the growth of *T. marneffei* in iron-deficient conditions.

**Figure 3.**
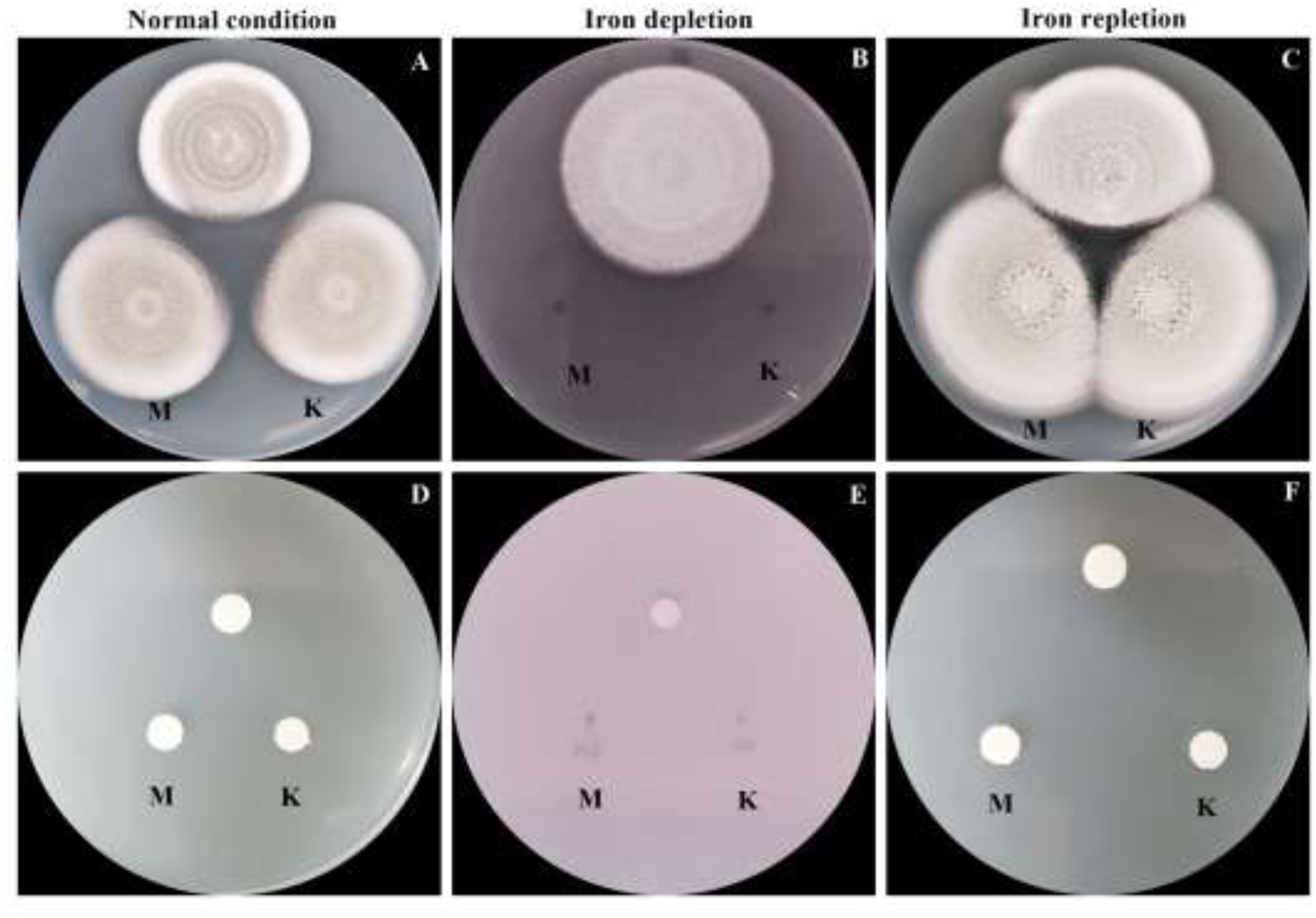
Growth of *T. marneffei* strains was determined in different iron conditions. Ten thousand conidia of the *T. marneffei* Δ*acuK*, Δ*acuM* and wild type ATCC18224 strains were spotted onto the surface of ANM agar. The media were prepared for the different iron conditions; normal medium (A and D ANM, 7 µM ferrous sulfate), iron-depletion medium (B and E ANM + 100 µM phenanthroline) and iron-repletion medium (C and F ANM + 100 µM phenanthroline and 1 mM FeCl_3_). The plates were incubated to grow as mold at 25°C (A – C) and as yeast at 37°C (D – F). Colony growth was monitored and photographed on day 12. M = *acuM* deletion mutant; K = *acuK* deletion mutant. Experiments were performed in three biological replicates.

### AcuK and AcuM affect the normal expression of genes involved in iron homeostasis

To determine if AcuK and AcuM controlled the expression of genes known to be involved in iron uptake, four gene candidates were selected that encode the iron transcription regulators *sreA* and *hapX*, and the RIA-enzymes *ftrA* and *fetC.* The upstream region of the selected genes was searched for the presence of the putative DNA binding motif of AcuK and AcuM (CCGN7CCG) and the results were listed in Table S2. Conidia of *T. marneffei* were cultured at 10^8^ conidia/ml in ANM broth supplemented with various iron concentrations (10 – 1000 uM of FeCl_3_). After growing at 25°C for 48 h, the cultures were harvested, and RNA samples were subjected to qRT-PCR.

As expected, the wild type exhibited increased levels of *sreA* gene expression and decreased levels of *hapX* gene expression as the iron concentrations were elevated from 10 to 1000 μM (Fig 4, A and B). However, *sreA* gene expression was highly upregulated in both mutants, especially under low iron conditions (Fig 4A). Furthermore, even though the mutants were able to downregulate *hapX* gene expression in response to different iron concentrations, the level of *hapX* gene expression was higher in the mutant strains when compared to the wild type across the tested conditions (Fig 4B). Remarkably, the *acuK* mutant exhibited higher *hapX* gene expression levels when compared to the *acuM* mutant in all tested iron concentrations (Fig 4B). These current results were consistent with our previous data reported on the *acuK* mutants (25). Thus, it seems that the *sreA*-*hapX* regulatory circuit is misregulated in the absence of AcuK or AcuM.

**Figure 4.**
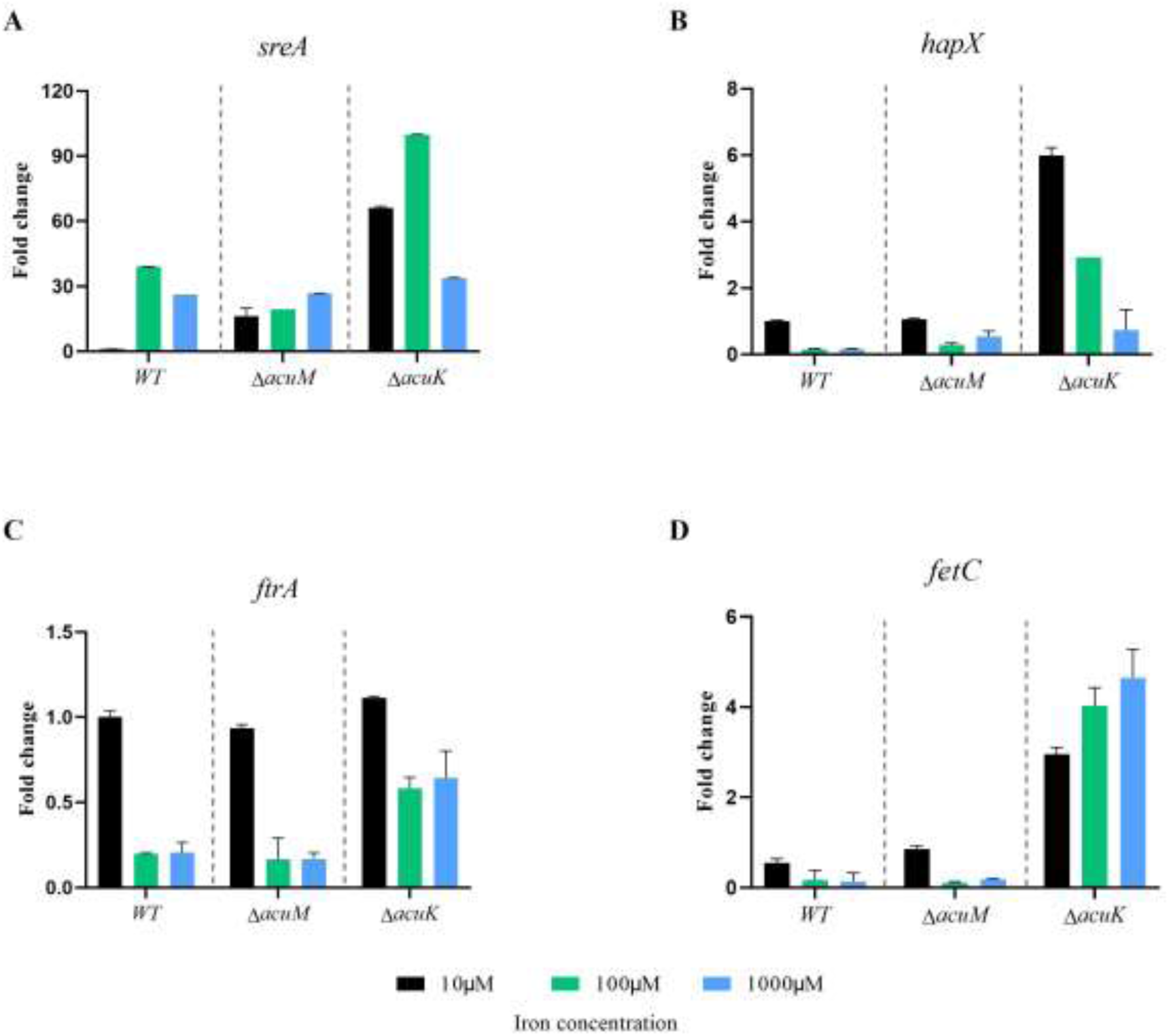
Expression analysis of iron uptake genes in the *T. marneffei* mutants under different iron concentrations. *T. marneffei* wild type, Δ*acuK* and Δ*acuM* strains were grown under various iron concentrations, in which 10, 100 and 1000 indicate micromolar concentrations of ferric chloride. The qRT-PCR was used to determine the fold-change of (A) *sreA*, (B) *hapX*, (C) *ftrA* and (D) *fetC* gene transcripts. Expression was compared to the wild-type strain under low iron concentration of 10 μM and the experiments were performed in triplicates.

For genes in the RIA system, the wild type strain downregulated the *ftrA* and *fetC* gene expression levels when iron concentrations were high to reduce iron toxicity as expected (Fig 4, C and D). The *acuM* mutant showed *ftrA* and *fetC* gene expression patterns that were similar to the wild type. However, the *acuK* mutants showed upregulated levels of *ftrA* gene expression at high iron levels and expressed the *fetC* gene at higher levels under all tested conditions (Fig 4, C and D). The high expression levels of genes encoding RIA enzymes in the *acuK* mutant was consistent with our previous study (25).

To investigate the involvement of *acuK* and *acuM* in siderophore biosynthesis, the amount of siderophore production was measured in the Δ*acuK* and Δ*acuM* mutants, using a Chrome Azurol S (CAS) solid and liquid assay. First, *T. marneffei* wild type and mutants were cultivated in ANM broth for 7 days. The amount of extracellular siderophores was measured from culture supernatant while the intracellular siderophores were determined from cell lysates. The production of intracellular and extracellular siderophores was unaffected in the Δ*acuK* and Δ*acuM* strains when compared to the wild type (Fig 5). This result demonstrates that the absence of either AcuK or AcuM does not affect siderophore production.

**Figure 5.**
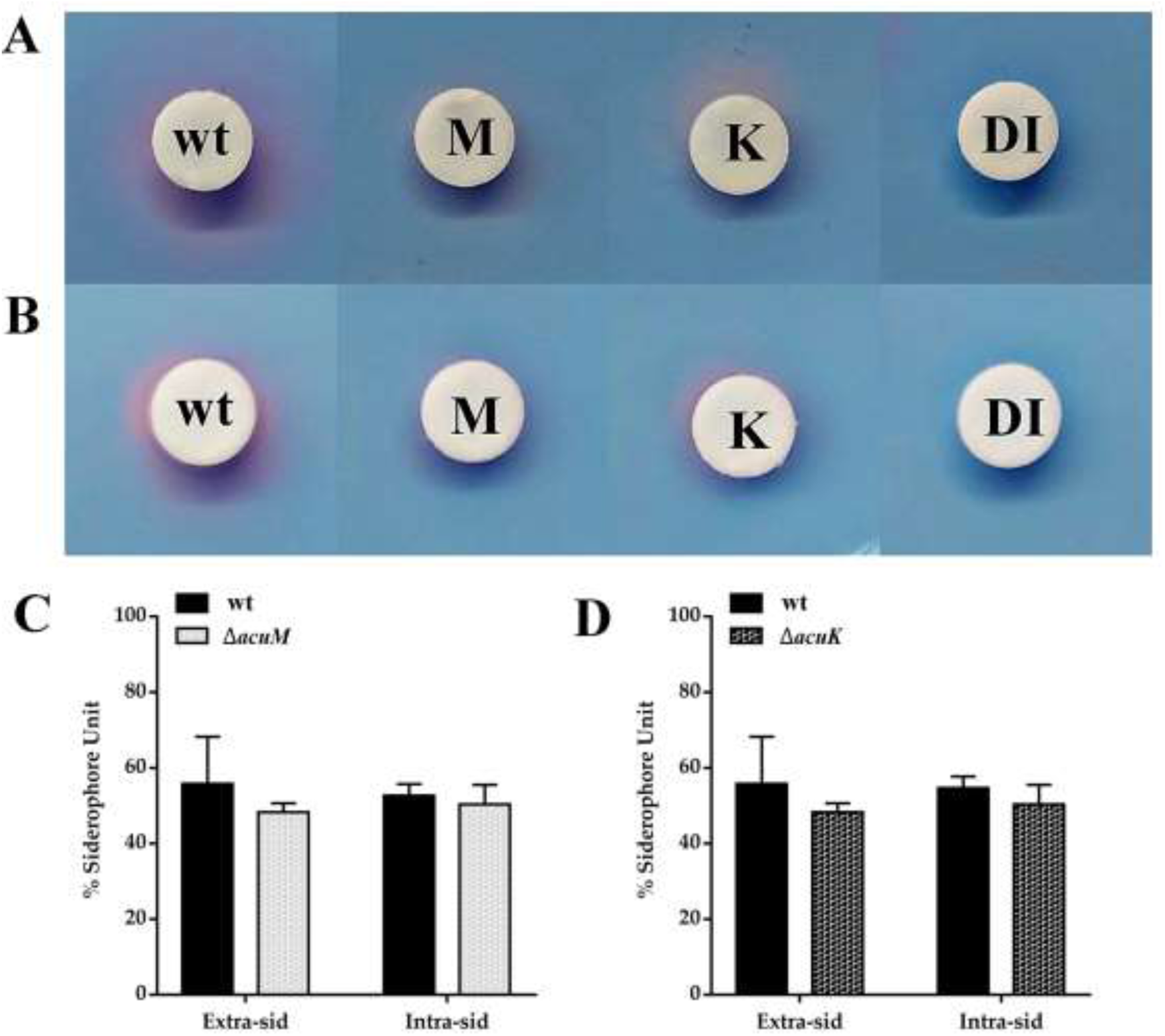
Siderophore production was determined in different *T. marneffei* strains. CAS agar diffusion (A and B) assay was performed to measure the siderophore generated by each *T. marneffei* strain. (A) Extracellular siderophore production was determined from culture supernatant of *T. marneffei* strains. (B) The cell lysate was used to determine the amount of intracellular siderophores. CAS liquid assay (C and D) was used for quantitative determination of the extracellular and intracellular siderophore production. *T. marneffei* strains included WT, wild type ATCC18224; K, Δ*acuK*; and M, Δ*acuM*. DI, distilled water served as a negative control. Error bars represent standard deviation of three experimental replicates.

Together, our results suggest that AcuK and AcuM might not directly activate genes in the RIA or siderophore-based iron acquisition pathways. Paradoxically, the iron acquisition genes were increased rather than decreased in the mutants. Hence, alternatively, other iron-dependent processes might lead to the impaired growth seen in Δ*acuK* and Δ*acuM* mutants under low iron conditions.

### Identification of AcuM and AcuK target genes

To gain insight into the molecular pathways dependent on AcuK and AcuM that have contributed to the observed phenotypes, we performed RNA-seq analysis and determined gene expression profiles in the Δ*acuK* and Δ*acuM* mutants versus the wild type strain. The experiments were only conducted in the hyphal growth phase due to the limited growth during the yeast phase in the mutant strains. To minimize growth differences between the mutant and wild type strains and to find global target genes under normal conditions, *T. marneffei* was grown in synthetic dextrose broth (1% glucose) at 25°C for 36 hours. The mRNA was then isolated and processed for RNA-seq analysis. The RNA-seq data were deposited at Gene Expression Omnibus (GEO) accession GSE248925.

For differential gene expression analysis, transcript levels (FKPM) of each mutant were compared to the reference wild type as described in materials and methods. The potential AcuK and AcuM target genes were defined as those genes in which expression was altered by a 2-fold or greater change with an adjusted P value of 0.05 or lower. The result identified 1,310 genes that were downregulated and 2,058 genes that were upregulated in the Δ*acuM* mutant vs the wild type. There were 1,286 downregulated genes and 2,214 upregulated genes in the Δ*acuK* mutant compared to the wild type. Transcriptome profiles of the Δ*acuK* and Δ*acuM* mutants showed a high correlation (Fig 6A, Pearson correlation coefficient > 0.95) and shared over 96% of differentially expressed genes (Fig 6B). Gene Ontology (GO term) and KEGG pathway analyses were performed on each of the AcuK-dependent genes and AcuM-dependent genes as listed in Table 1.

**Figure 6.**
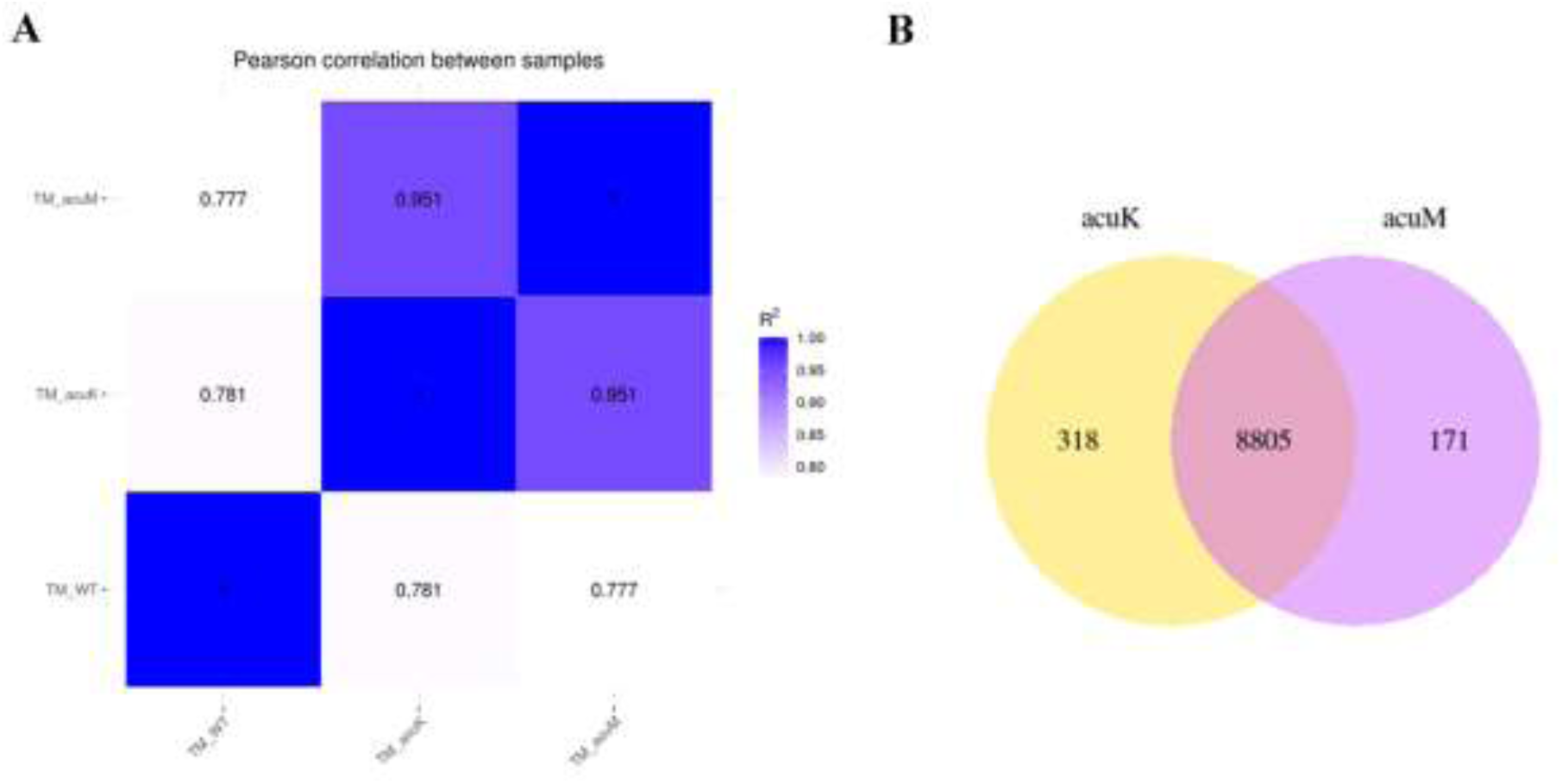
Transcriptome profiles of the *acuK* and *acuM* mutants were analyzed by RNA-seq experiment. RNA-seq was performed in mold phase as described in materials and methods. (A) Heatmap depicted Pearson correlation coefficient (R^2^) between wild type, Δ*acuK*, and Δ*acuM* strains. (B) Venn diagram presented the number of DEGs that were uniquely and commonly expressed between the two mutants.

**Table 1.**
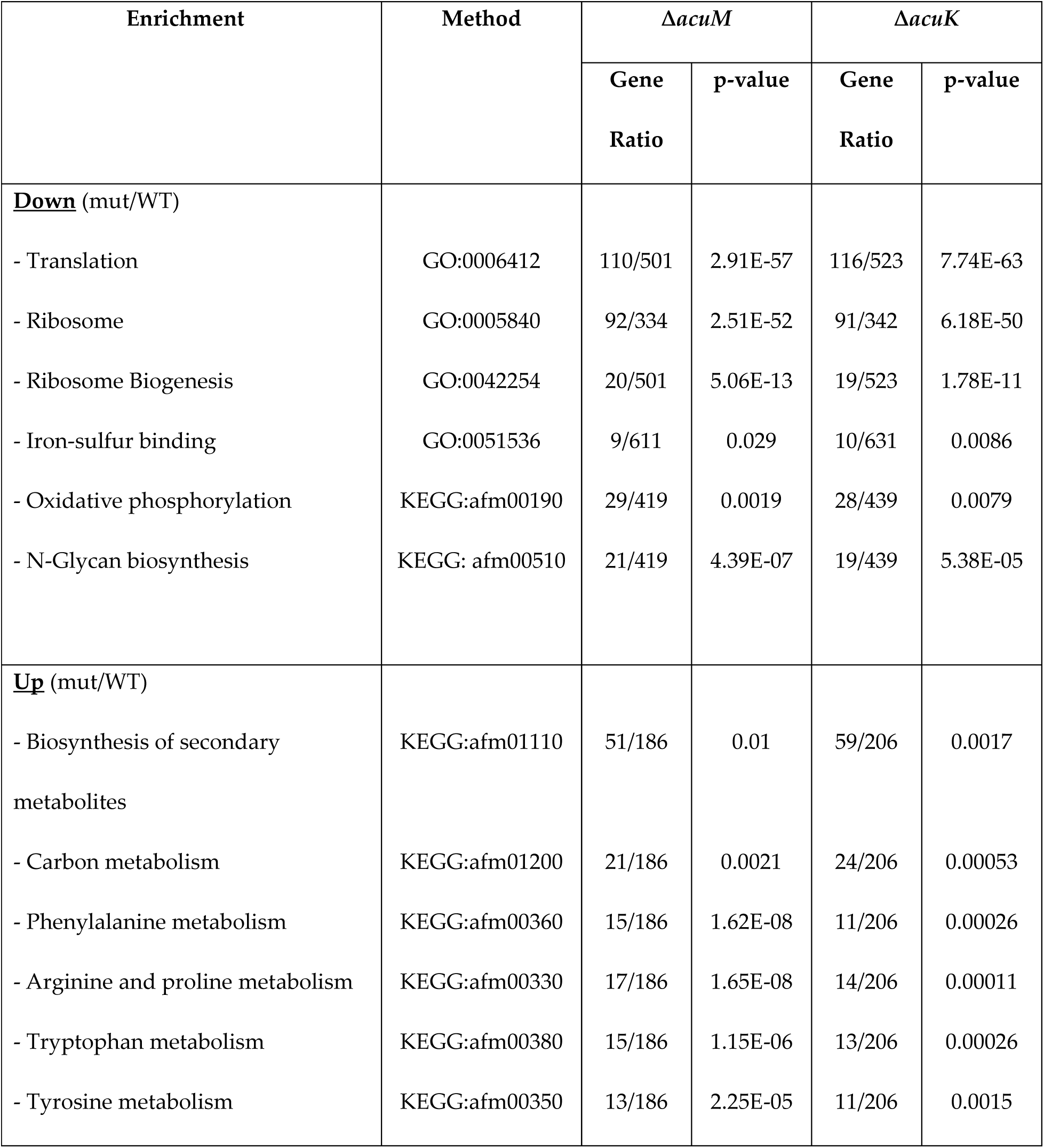
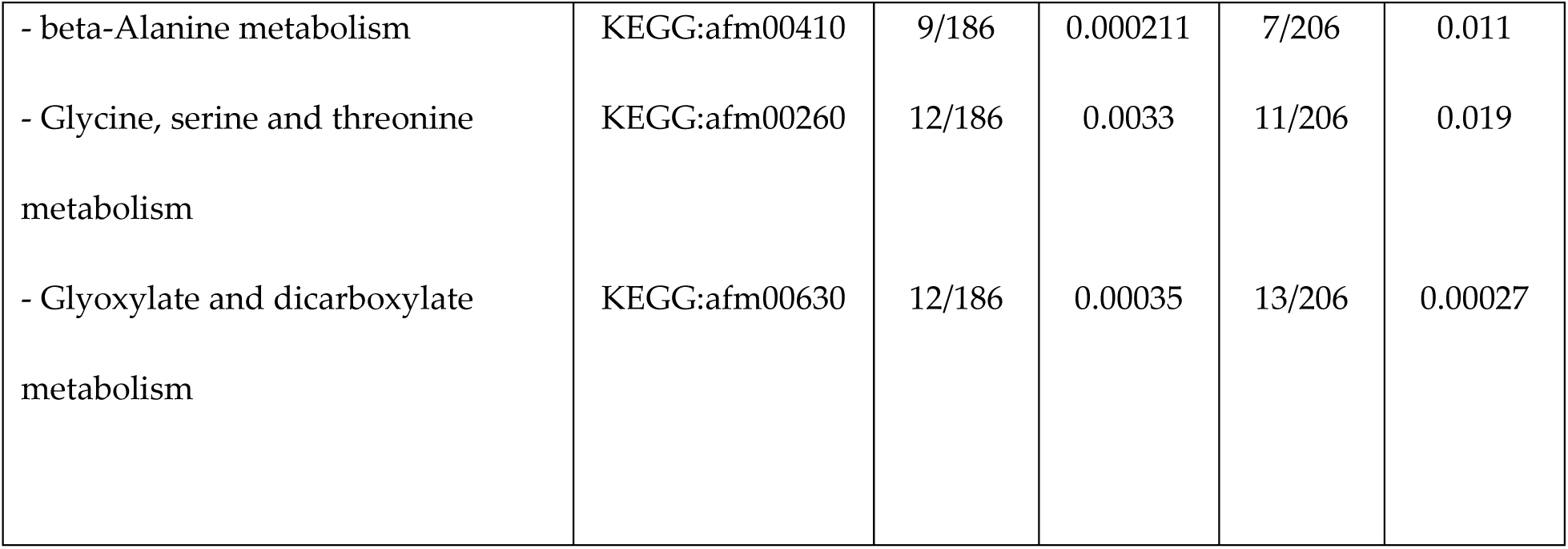
Overrepresented functional enrichments of *T. marneffei* genes in the Δ*acuK* and Δ*acu*M mutants vs wild type (log_2_fold-change ≤ −1 or ≥ 1, and p-value ≤ 0.05).

GO term and KEGG analyses indicated that the downregulated genes in the Δ*acuK* and Δ*acuM* mutants were linked to oxidative phosphorylation and iron-cluster binding (Table 1 and Fig 7). For instance, the cytochrome C gene (PMAA_076160) was expressed at an 8-fold lower level in the mutants when compared to the wild type (Table S3). The alternative respiration enzyme, iron-containing alternative oxidase (AOX, PMAA_029240), was downregulated approximately 2-fold in the Δ*acuK* and Δ*acuM* strains (Table S3). The expression of alternative NADH dehydrogenase (PMAA_065210), another alternative respiration enzyme, was expressed at lower levels in both mutants, exhibiting a 20-fold and 11-fold decrease in the *ΔacuM* and *ΔacuK* strains, respectively (Table S3). This prompted us to inspect the expression of genes involved in other iron-consuming pathways such as iron-sulfur cluster biosynthesis proteins, iron-regulated transporters, and iron-containing proteins of the TCA cycle. Genes in these pathways were significantly downregulated in the Δ*acuK* and Δ*acuM* mutants (Fig 8). For example, the gene encoding for succinate dehydrogenase (PMAA_089220) was downregulated by 4-fold in the Δ*acuK* and Δ*acuM* strains (Table S3). Moreover, several genes encoding for iron-containing proteins that participate in the antioxidative stress response were differentially expressed in the mutants. For example, the iron superoxide dismutase (FeSOD, PMAA_058080) gene was downregulated approximately 3-fold in the Δ*acuM* and Δ*acuK* strains (Table S4). Interestingly, the mycelial catalase gene (*CAT1*, PMAA_014760) was expressed at higher levels in the mutants, being upregulated 4-fold in each mutant (Table 3). The *CAT2* gene encoding for catalase-peroxidase enzyme (CPE) was expressed 4-time higher in the Δ*acuK* mutant (Table S4). In addition, ribosomes and ribosome biosynthesis genes were enriched in the downregulated gene set of both the Δ*acuK* and Δ*acuM* mutants (Fig 7, Table 1 and S5). This result suggests the Δ*acuK* and Δ*acuM* mutants might have impaired the protein translation process. Overall, our data showed that while AcuK and AcuM did not directly regulate genes involved with iron acquisition, genes involved in iron-consuming processes were targets. Because multiple pathways utilize iron, defective expression of these iron-dependent proteins could affect the growth of the mutants under various harsh environmental conditions. Based on the transcriptomic profile, AcuK and AcuM seemed to control a broader aspect of mitochondrial metabolism than anticipated.

**Figure 7.**
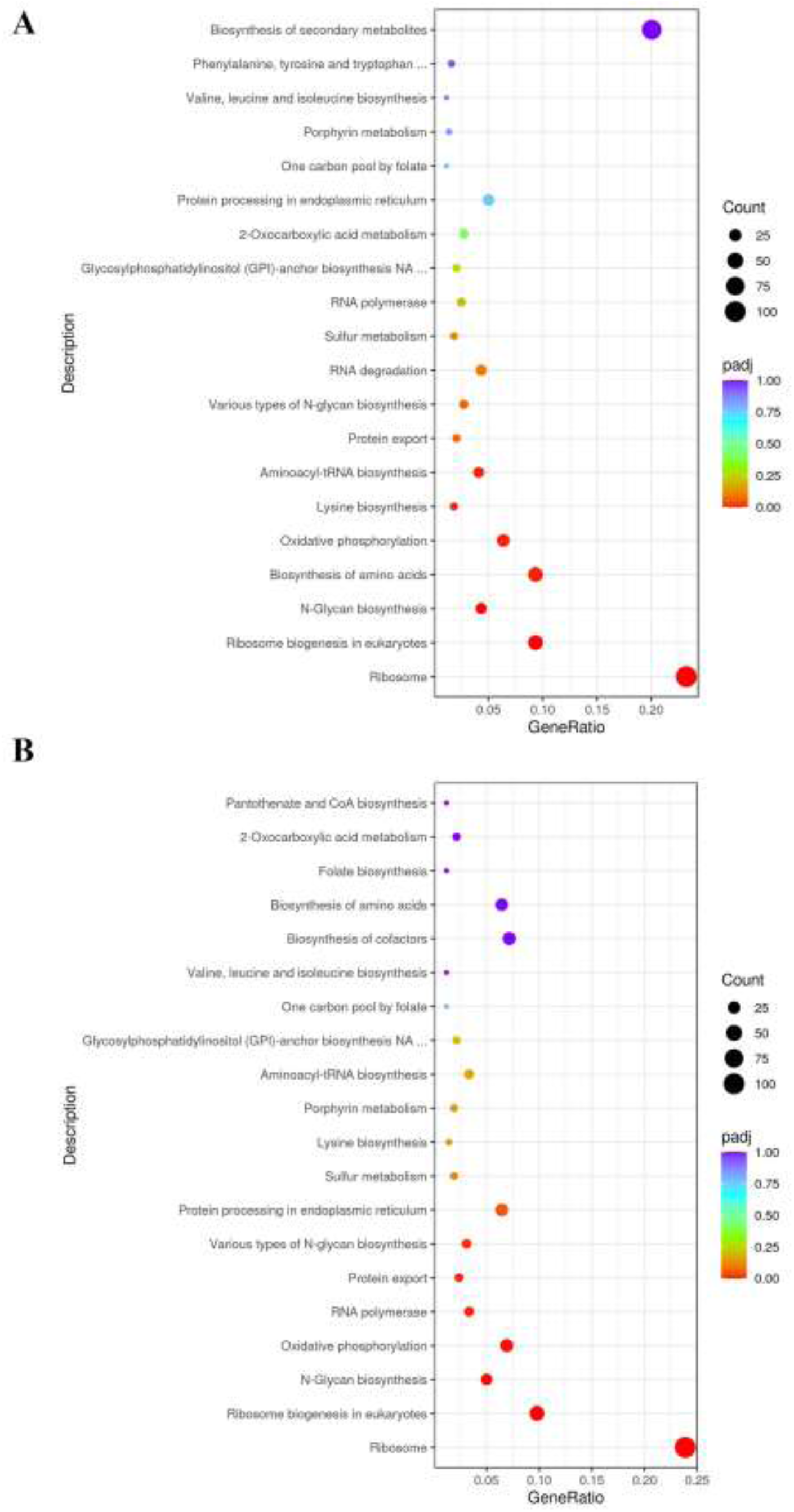
KEGG analysis depicted downregulated pathways that were enriched in the *acuK* and *acuM* deletion mutants compared to the wild type. Enrichment analysis of differentially downregulated genes in in the (A) Δ*acuM,* and (B) Δ*acuK* deletion mutants was performed as described in materials and methods.

**Figure 8.**
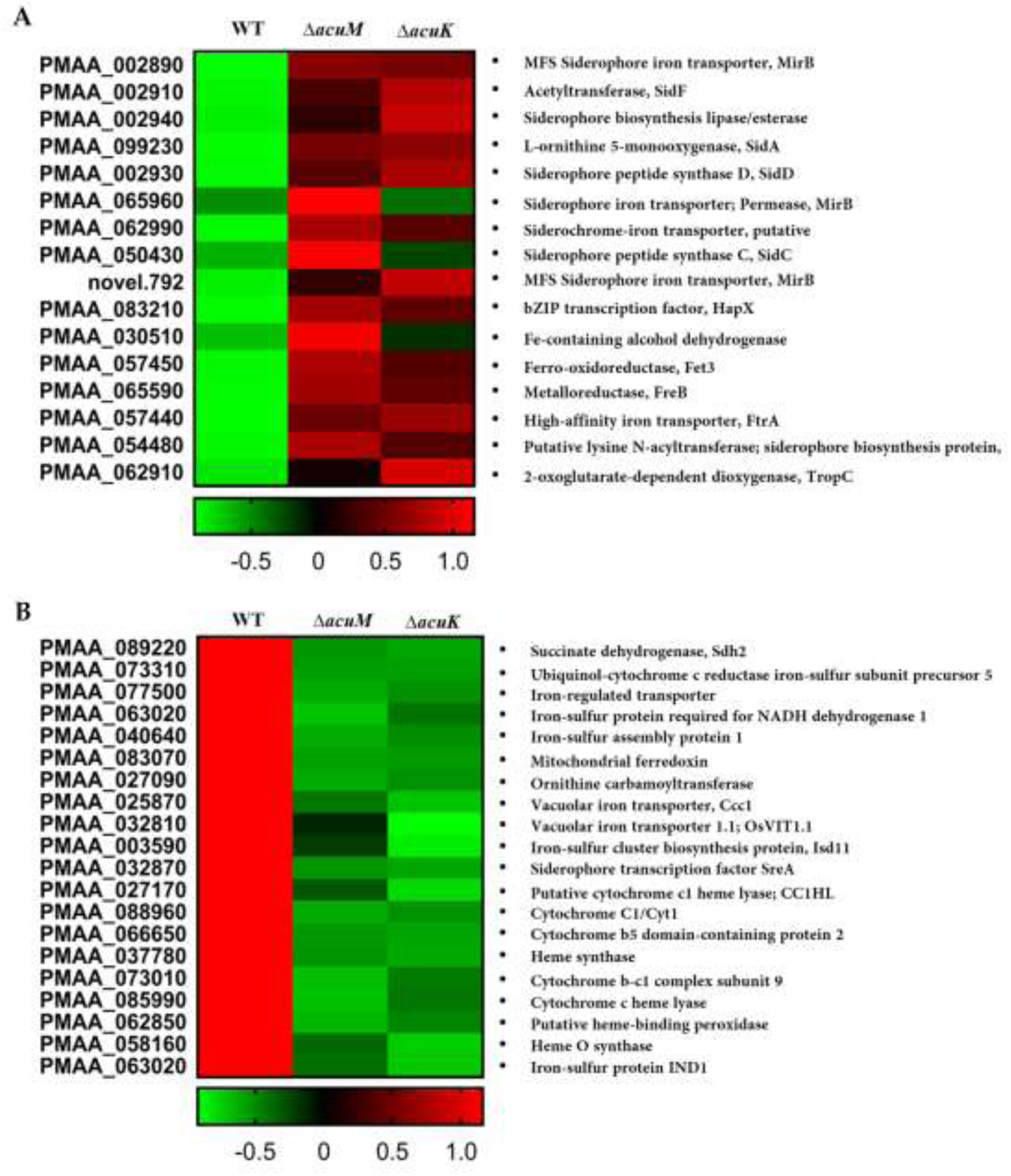
Heatmap represented genes involved in iron metabolism that were differentially expressed in the mutants compared to wild type. Genes of interest were manually selected from transcriptome data, and their expression levels (log_2_(FPKM+1) values) were depicted as a Heatmap. (A) heatmap of gene group in iron acquisition pathway. (B) heatmap of gene groups in iron consumption pathway. Heat map score: Red indicates higher expression levels; Green indicates lower expression levels compared to the wild type.

Next, we manually determined if the known genes specific to gluconeogenesis and iron homeostasis were differentially expressed in the mutants under tested conditions (Table S4). The transcriptome data revealed that Δ*acuK* and Δ*acuM* strains were unable to induce the expression of *acuF,* even in the presence of glucose (Table S4). This *acuF* gene encodes for phosphoenolpyruvate carboxykinase (PEPCK), another known key gluconeogenic enzyme regulated by AcuK and AcuM homologs (17). By combining this information with the qRT-PCR data (Fig 2), it suggests that AcuK and AcuM of *T. marneffei* could activate gluconeogenesis through the transcriptional regulation of *fbpA* (*acuG*) and *acuF* genes. Also, iron acquisition genes encoding for RIA proteins and siderophore synthetic enzymes were expressed at significantly higher levels in the Δ*acuK* and Δ*acuM* mutants (Table S4 and Fig 8A), consistent with our qRT-PCR gene expression studies (Fig 4 and (25)). We noted that the higher expression levels of multiple iron acquisition genes did not lead to a higher amount of siderophore production in both mutants (Fig 5). Overall, AcuK and AcuM could affect the expression of gluconeogenic and iron homeostasis genes even under non-inducing conditions (i.e. in the presence of glucose and iron).

### AcuK and AcuM are necessary for oxidative stress response

In addition to the function of respiration, alternative oxidase and alternative NADH dehydrogenase enzymes could participate in antioxidant defenses against oxidative stress (31, 32). As the expression of these genes was significantly downregulated in the Δ*acuK* and Δ*acuM* mutants (Table S3 and S4), we hypothesized that the two mutants could not cope with oxidative stress. During normal metabolic processes, organisms can produce a range of ROS, including superoxide, the hydroxyl radical, and hydrogen peroxide, and the different types of ROS can lead to different metabolic and transcriptomic responses (33). Accordingly, we examined the growth of the Δa*cuK* and Δ*acuM* mutants in ANM containing menadione (to generate intracellular O_2_^•−^) at 5 µM and 10 µM and exogenous H_2_O_2_ at 1-5 mM.

The Δ*acuM* mutant showed growth impairment to menadione (Fig 9A) and hydrogen peroxide (Fig 9B) treatment. The growth sensitivity was more evident in the yeast phase than the mold phase (Fig 9). On the other hand, the growth of the Δa*cuK* mutant was not affected by menadione (Fig 9A) while the mutant exhibited growth sensitivity to hydrogen peroxide treatment (Fig 9B). This result prompted us to examine the expression levels of the anti-stress genes from transcriptomic data. Importantly, several genes with antioxidant roles (*SOD*, *CAT1,* and *CAT2 (CPE))* showed differential gene expression in the Δ*acuM* and Δ*acuK* mutants (Table S4). Together, these findings indicate that AcuM plays a more prominent role in response to oxidative stress than AcuK. Also, the variations in phenotypes between the two mutants suggest that AcuK and AcuM can regulate some target pathways independently.

**Figure 9.**
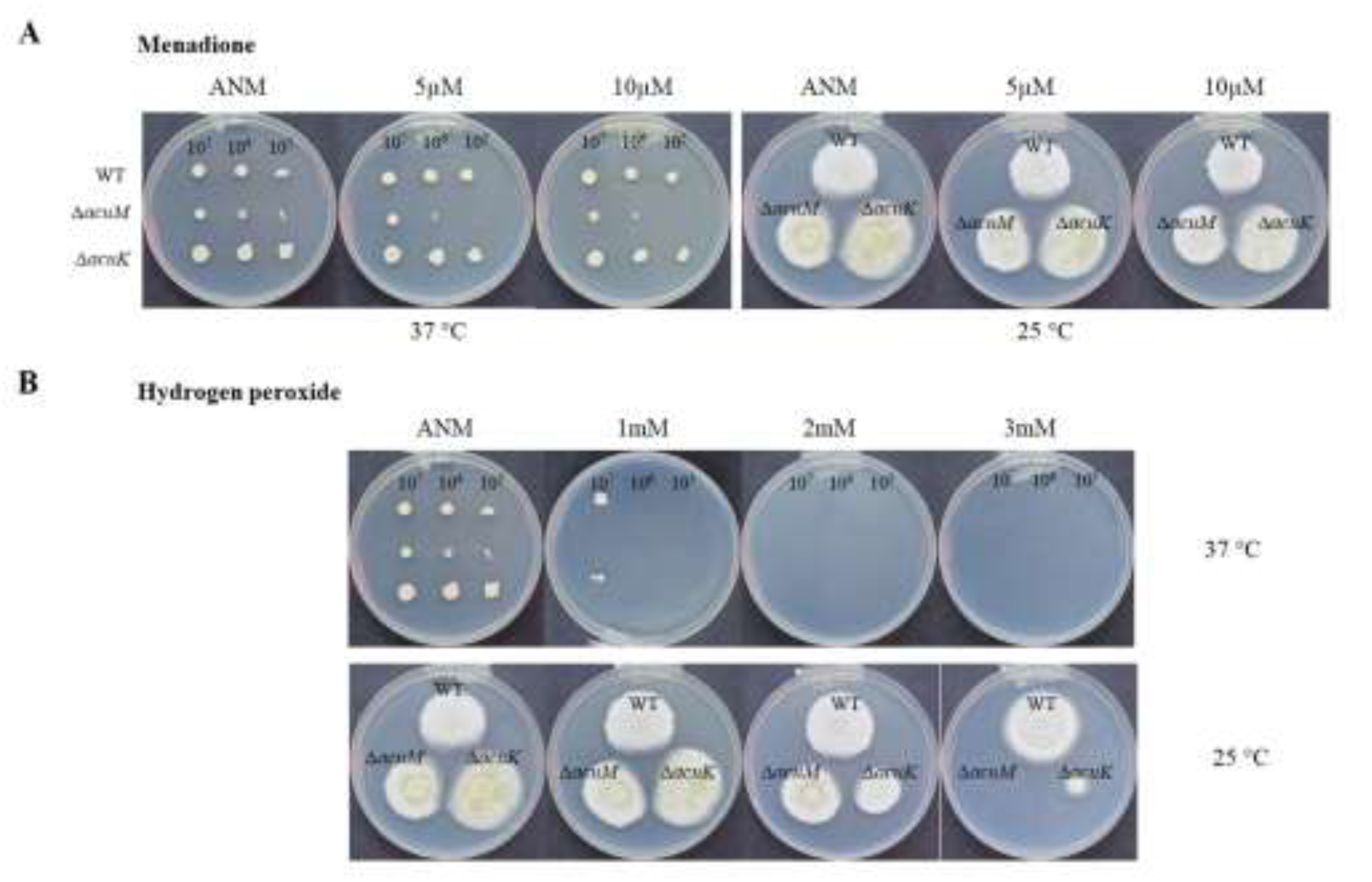
Growth of the *acuK* and *acuM* deletion mutants were examined in the presence of oxidative stressors. *T. marneffei* wild type, Δ*acuK* and Δ*acuM* strains were grown in ANM medium containing (A) menadione, and (B) H_2_O_2_ at indicated concentrations. They were incubated at 25 °C and 37 °C. Photographs were taken on day 14. Experiments were conducted in three replicates.

### Bioinformatics analysis of binding sites of AcuK and AcuM

The DNA binding motif of AcuK and AcuM contains repeats of CGG separated by seven bases (CCGN_7_CCG) (17). This motif is conserved across *Aspergillus* species, and it is commonly found in the promoter of genes coding for enzymes in the TCA cycle, glycolysis, and gluconeogenesis. To identify potential target genes for AcuK and AcuM in *T. marneffei,* the upstream region of genes in the genome were searched for putative CCGN_7_CCG consensus binding sites using the web-based program FIMO (Find Individual Motif Occurrences) as described in materials and methods (Table S2). The CCGN7CCG sequences were present in many genes that were differentially regulated in the mutants from the RNA-seq data, including genes involved with the TCA cycle, glycolysis, and ribosome biogenesis (Table S2). Consistently, GO analysis of identified genes that contain the CCGN_7_CCG sequences at the upstream region revealed enrichment in transcriptional regulation, mitochondrial functioning (proteins encoded by or localized in the mitochondrion), and heme/iron binding (iron-dependent proteins). Overall, our data suggests that AcuK and AcuM could potentially bind directly to the promoters of these target genes and regulate their expression.

### AcuM is necessary during macrophage infection

To assess if *acuM* affected *T. marneffei* interactions with host cells, differentiated THP-1 macrophage cells were infected with conidia from wild type G809 (*acuM*^+^) and Δ*acuM* strains. After 24-hr of infection, the *T. marneffei* wild type strain produced short filaments within the THP-1 macrophages (Fig 10). In contrast, only non-germinated cells were found in the Δ*acuM* mutant-infected cells. The percentage of phagocytosis, the phagocytosis index, and percentage of killing were determined at 2 and 24 hours of infection. As shown in the result (Table 2), the percentage of phagocytosis was decreased by 2-fold and 3-fold in the Δ*acuM* mutant compared to the wild type at 2- and 24-hour time points, respectively. However, the percentage of killing in the mutant was slightly increased, only 11% higher than the wild type (Table 2). This result together implied that AcuM is important for host-pathogen interaction, especially during macrophage engulfment.

**Figure 10.**
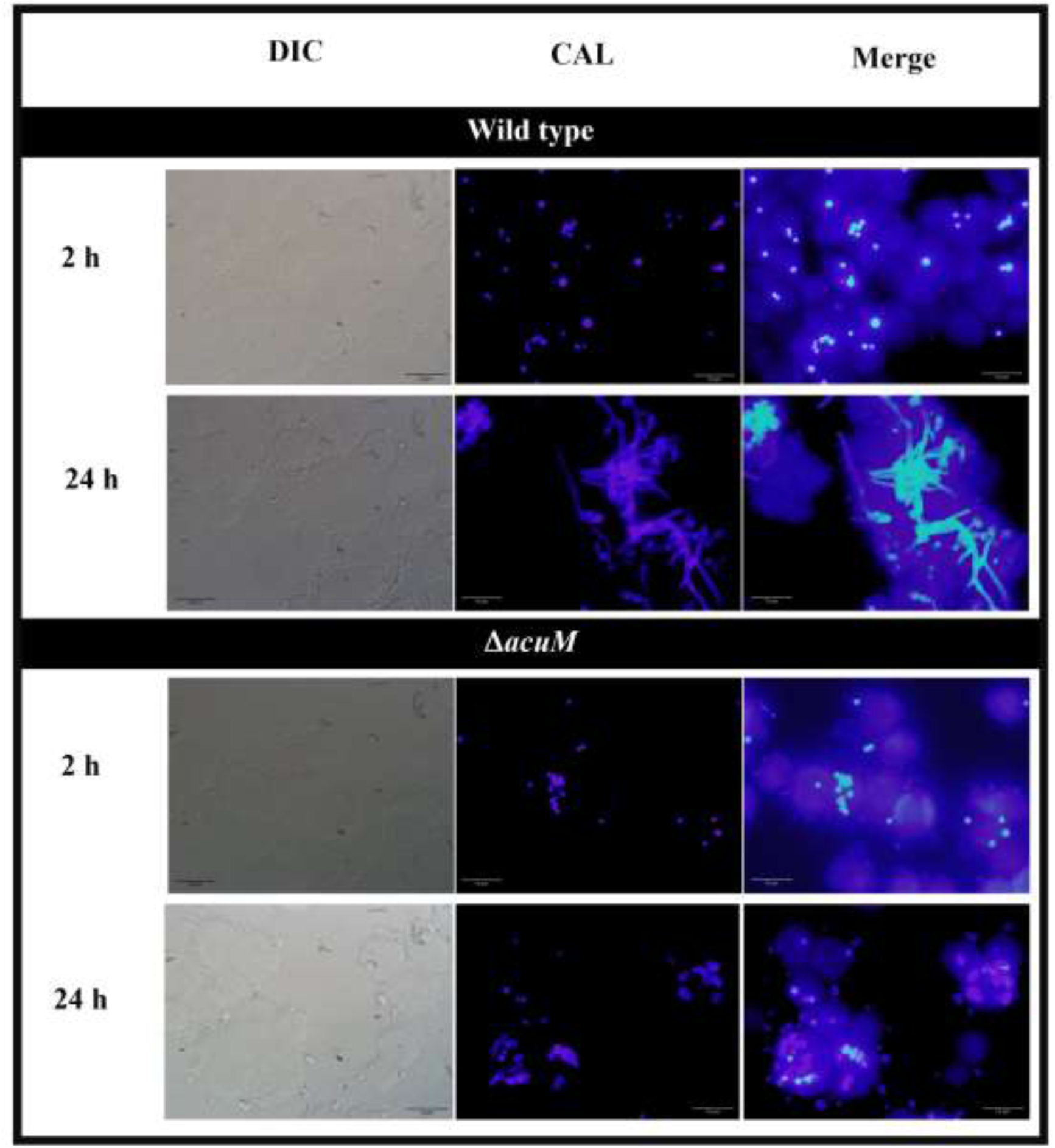
The calcofluor white-labeled *T. marneffei* strains were examined in the THP-1 human macrophage infection model. PMA-activated THP-1 human macrophages were infected with conidia from wild type and Δ*acuM* strains and microscopically evaluated at 2- and 24-hr (scale bars are 10 μM, 100X magnification). Experiments were conducted in three replicates.

**Table 2.** Percentage of phagocytosis, phagocytic index, and percentage of killing of *T. marneffei* strains by the macrophage THP-1.

| Relevant genotype | Incubation time (hr) | % Phagocytosis | Phagocytic index | %Killing |
| --- | --- | --- | --- | --- |
| *acuM*<sup>+</sup> | 2 | 40 | 1.43 | 66.12 |
|  | 24 | ND | ND | 99.64 |
| Δ*acuM* | 2 | 18 | 2.06 | 73.58 |
|  | 24 | ND | ND | 99.97 |

ND = Not determined

## DISCUSSION

Our current data suggests a larger role of the AcuK and AcuM proteins as global regulators of cell growth and metabolism in *T. marneffei*. First, we demonstrated that AcuK and AcuM play an important role in regulating gluconeogenesis. Neither Δ*acuK* nor Δ*acuM* strains of *T. marneffei* were able to grow on the tested non-glucose carbon substrates ((25) and this study). Our gene expression analyses showed that Δ*acuK* and Δ*acuM* strains were unable to induce the expression of *fbpA* (*acuG*, data by qRT-PCR) and PEPCK (*acuF*, data by RNA-seq) genes, which encode the two known key gluconeogenic enzymes. The role of *T. marneffei* AcuK and AcuM in regulating gluconeogenesis via transcriptional control of *acuG* and *acuF* gene expression is highly conserved in other fungal species, including *A. nidulans* (17)*, A. fumigatus* (20, 21)*, Neurospora crassa aod2* and *aod5* (34), and *Podospora anserina rse2* and *rse3* (35).

Second, in addition to gluconeogenic genes, our transcriptome data demonstrated that AcuK and AcuM positively regulate the expression of genes encoding for alternative oxidase, alternative NADH dehydrogenase, cytochrome C protein, and malic enzyme. Indeed, the genome-wide studies in *N. crassa* and *P. anserine* revealed that the AcuK and AcuM homologs commonly regulated the same set of target genes (34–37). Among these identified target genes, the transcriptional induction of the alternative oxidase gene by *N. crassa* aod2/5 and *P. anserina* rse2/3 transcription factors have been well characterized. Alternative respiration pathways allow electron transport and continuity of respiration in the presence of mitochondrial respiration inhibitors such as antimycin A and cyanide. In the context of fungal infection, nitric oxide, generally produced from host defense mechanisms, is a potent inhibitor of mitochondrial respiration, yet it has no inhibitory effect on alternative oxidase. (38, 39). The studies in *A. nidulans*, *N. crassa* and *P. anserina* suggested that the expression of gluconeogenic and alternative oxidase genes by AcuK and AcuM proteins is induced by different environmental signals (i.e. alternative carbon source availability vs mitochondrial respiration impairment). Overall, the role of *T. marneffei* AcuK and AcuM in regulating gluconeogenic genes and alternative respiration component genes is highly conserved in other fungal species.

Third, we investigated the role of AcuM and AcuK in the regulation of iron homeostasis. Interestingly, the role of AcuK and AcuM transcription factors in response to iron availability seems to be restricted to only pathogenic fungi. To date, this role has not been reported in the non-pathogenic mold *A. nidulans*, *N. crassa*, and *P. anserina*. At the phenotype level, Δ*acuK* and Δ*acuM* strains of *T. marneffei* could not grow on iron-depleted medium. This similar phenotype is observed in the Δ*acuK* and Δ*acuM* strains of *A. fumigatus* (20, 21). However, the molecular basis underlying the observed phenotype is drastically different between *A. fumigatus* and *T. marneffei*. In *A. fumigatus*, AcuK and AcuM activate gene expression of iron uptake genes. Deletion of *acuK* and *acuM* genes in *A. fumigatus* leads to downregulation of the *hapX* gene as well as genes in the RIA and siderophore production systems, which ultimately leads to a decrease in iron incorporation and extracellular siderophore production (20, 21). As opposed to the gene expression patterns detected in *A. fumigatus*, deletion of *acuK* and *acuM* in *T. marneffei* increased the expression of both the *hapX* gene as well as the RIA and siderophore biosynthesis genes under low, normal, and high iron conditions. It is possible that carbon metabolic gene disruption occurring in the Δ*acuK* and Δ*acuM* strains could lead to a cellular response similar to iron starvation. This result indicates that AcuK and AcuM from *T. marneffei* were not directly involved with the activation of the iron acquisition system as reported in *A. fumigatus*. Instead, our transcriptome analysis revealed that the expression of gene-encoded proteins containing iron-sulfur clusters and those participating in oxidative phosphorylation were strongly downregulated in the Δ*acuK* or Δ*acuM* strains. Thus, the impaired growth noted in the *T. marneffei* Δ*acuK* and Δ*acuM* mutants under low iron conditions could result from the inability to utilize iron-dependent proteins, rather than the inability to scavenge iron from the environment as seen in *A. fumigatus*.

How could iron-utilization defects disrupt the growth of the Δ*acuK* and Δ*acuM* strains in *T. marneffei*? Considering that multiple proteins of cellular respiration utilize iron (40, 41), the iron-utilization defects in the *T. marneffei* Δ*acuK* and Δ*acuM* mutants could disrupt many essential metabolic pathways that supply metabolites and energy to the cells. Indeed, it has been previously reported that mitochondrial metabolism (i.e., the TCA cycle and oxidative metabolism) plays a greater role in the pathogenic yeast form of *T. marneffei* (14). Inactivation of the TCA cycle completely prohibits the growth of *T. marneffei* yeast cells (14). This is consistent with the fact that there are more mitochondria in the yeast form compared to the filamentous form of *T. marneffei* (8, 42). Furthermore, the yeast form is more efficient in metabolizing a diverse range of carbon sources when compared to the mycelium form (14). Importantly, the ability of fungi to utilize various carbon sources metabolized through the TCA cycle requires gluconeogenesis, implying that gluconeogenesis is more important in the *T. marneffei* yeast form compared to the mold form (17). This phase-specific metabolic profile is consistent with our result that, yeast cells lacking *acuK* and *acuM* displayed more severe growth defects than the hyphal cells under both low iron and gluconeogenic conditions. Although the connection between gluconeogenesis and alternative respiration is not obvious, our findings suggests that the regulation of carbon metabolism and iron homeostasis is strongly connected. Overall, iron-utilization defects, especially in enzymes participating in both classical and alternative respiration pathways, could disrupt growth in the Δ*acuK* and Δ*acuM* strains. Reciprocally, the inability of the Δ*acuK* and Δ*acuM* strains to utilize non-glucose carbon sources could be a result of defects in multiple iron-dependent enzymes beyond the gluconeogenic enzymes.

Fourth, our transcriptome data revealed for the first time that AcuK and AcuM are implicated in the transcriptional regulation of ribosome biogenesis and protein translation. The reduction of protein synthesis components could halt the overall growth in the Δ*acuK* and Δ*acuM* mutants, contributing to the growth defects seen in the mutants under gluconeogenic and iron-insufficient conditions. In fact, high transcript levels of RIA genes, and siderophore biosynthesis genes observed in the mutants may not get translated into final protein products. Translation inhibition is a general response to several stresses, including osmotic, oxidative, stress, heat shock, and nutritional deficiencies of glucose and amino acids (43–47). Recently, it has been shown in *S. cerevisiae* that a global translation arrest occurs under iron deficiency. Many steps in the translation process utilize iron-containing enzymes, including ribosome biogenesis and recycling, translation initiation and termination, and modification of translation elongation factors and transfer RNAs (tRNAs). For instance, the iron-sulfur protein Rli1 is required for translation initiation, translation termination, and ribosome biogenesis/recycling (48–52). In budding yeast *S. cerevisiae*, TORC1 and Gcn2/eIF2a pathways regulate global translation repression (53) while the post-translation regulator Cth2 mediates specific translation repression of iron-utilizing proteins (43, 54–56). To our knowledge, AcuK and AcuM are the first potential set of transcription factors reported in fungal pathogens that coordinate the expression of ribosome biogenesis genes with the expression of iron homeostasis genes. Future investigations will decipher how AcuK and AcuM regulate the translation process and response to nutrient availability.

Also, we demonstrated that AcuK and AcuM from *T. marneffei* regulated the response to oxidative stressors. The activity of alternative respiration components is known to be associated with metabolic homeostasis, ROS control, and stress response in fungi (32, 38, 57). This is because alternative oxidase and alternative NADH dehydrogenase are non-proton pumping proteins that can bypass several steps of the electron transport chain in mitochondria, and therefore alternative respiration pathways could lower endogenous ROS levels. The Δ*acuM* mutant exhibited growth sensitivity to intracellular superoxide generated by menadione and to extracellular hydrogen peroxide. Thus, a decrease in genes encoding for superoxide dismutase, alternative oxidase, and alternative NADH dehydrogenase in the Δ*acuM* mutant likely leads to sensitivity to oxidative stressors. In contrast to the Δ*acuM* mutant, the Δ*acuK* mutant showed normal growth under menadione treatment and only mild growth sensitivity to hydrogen peroxide, suggesting that oxidative stress response is less dependent on AcuK. Consistent with the phenotype, upregulated levels of catalase-peroxidase *CAT2 (CPE)* gene were detected only in the Δ*acuK* mutant. Thus, we postulate that high expression levels of the catalase-peroxidase gene may compensate for downregulated levels of superoxide dismutase, alternative oxidase, and alternative NADH dehydrogenase genes, contributing to the normal growth observed in the Δ*acuK* mutant. It is noteworthy to mention that the role of AcuK and AcuM in protecting cells against ROS via the regulation of alternative respiration pathways has never been investigated in any fungal species. Mutations in homologs of *acuK* and *acuM* in *A. nidulan*s (17), *N. crassa* (36), and *P. anserina* (37) lead to growth sensitivity to antimycin A due to an inability to induce gene encoding alternative oxidase. Importantly the use of antimycin A could exacerbate the formation of mitochondrial ROS (31). The recent study by Emri et al. reveals that oxidative stress response in *A. fumigatus* is highly dependent on glucose and iron availability (62). Siderophore production and synthesis of Fe-S cluster proteins are crucial for cells to cope with iron limitation and oxidative stress (62). The expression of the alternative oxidase *aoxA* gene is dependent on iron availability and the presence of H_2_O_2_, suggesting the role of *aoxA* in maintaining the redox status of mitochondria during the oxidative stress response (62). Based on our results, we speculate that the AcuK/AcuM protein family likely participates in antioxidant mechanisms as well as other combinatorial stress responses in multiple fungi.

The macrophage killing assay of Δ*acuK* and Δ*acuM* showed similar results (25) and this study). Both mutants demonstrated an approximately 10% higher macrophage killing by THP-1 macrophage cells when compared to the wild type. This result implicates that the inactivation of the *acuK* and *acuM* genes could have enhanced fungal clearance by the host macrophage. Unlike the *T. marneffei* Δ*acuK* mutant (25), phagocytosis was decreased by over 2-fold in the Δ*acuM* mutant when compared to wild type. This result suggests that AcuM contributes to the ability of *T. marneffei* to interact with macrophage cells. One possibility could involve the alteration of glycoprotein synthesis that results in the rearrangement of cell wall components. Glycoproteins are components of the fungal cell wall and plasma membrane, which are the first points of contact with the host, and hence glycoproteins contribute to fungal pathogenicity, virulence, and host immune response. In this study, genes in N-glycan biosynthesis were downregulated in the Δ*acuM* mutant (Table 1). N-glycans can alternate the properties of glycoproteins including their activity, antigenicity and recognition by glycan-binding proteins (58). Accordingly, we postulate that changes in N-glycan biosynthesis in the Δ*acuM* mutant could be a factor in the alteration of cell wall and plasma membrane composition, contributing to a decrease in phagocytosis by THP-1 macrophage cells. The relevance of fungal protein glycosylation in host-pathogen interaction and virulence has been demonstrated in many medically important fungi (59). The N-glycan biosynthetic genes are also downregulated in the Δ*acuK* mutant. It is possible that the defects in N-glycan biosynthetic gene expression in the Δ*acuK* mutant may not result in functional defects as seen in the case of the Δ*acuM* mutant. This is consistent with the general observation that the Δ*acuM* mutant exhibits more severe phenotypic defects than the Δ*acuK* mutant. Alternatively, there might be other unknown relevant factors that could affect the phenotype of Δ*acuM*. In addition, the homologs of AcuM from *A. fumigatus* are required for maximal virulence in animal models of infections (20). It remains to be elucidated whether or not the absence of the *acuM* gene affects the virulence of *T. marneffei* as demonstrated in *A. fumigatus* by the mouse model of infection. Overall, the AcuK/AcuM represents a protein family that is important for host-pathogen interaction and virulence in several fungal pathogens.

As proposed by Bovier et al, the *rse2* and *rse3* transcription factors are involved with adaptation and defense of organisms to the environment (35). For intracellular fungal pathogens, they must adapt to metabolize alternative carbon sources (60, 61) available in the infected sites and defend against the host’s nutritional immunity, and host-derived oxidative stress (10). In *T. marneffei*, AcuK, and AcuM regulate the expression of target genes involved in at least four different pathways, including alternative carbon utilization, alternative respiration, iron homeostasis, and oxidative stress response. Thus, the AcuK and AcuM proteins likely are important for the virulence of *T. marneffei*. The essentiality of heterodimerization of two proteins for target gene activation has been confirmed in *A. nidulans* and *N. crassa* (17, 36). On one hand, the Δ*acuK* and Δ*acuM* mutants exhibit similar gene expression profiles and phenotypes in response to gluconeogenic substrates, and therefore we speculate that AcuK/AcuM could form a heterodimer to control the expression of gluconeogenic genes. On the other hand, there are differences in gene expression patterns and phenotypes in response to iron availability and oxidative stress, suggesting that AcuK and AcuM can regulate divergent targets as homodimers. In summary, this work demonstrates that AcuK and AcuM control multiple fitness attributes that are important for pathogenicity of *T. marneffei* during infection.

## AUTHOR CONTRIBUTIONS

**Conceptualization**: MP

**Data Curation**: AAm, PS, TK, JJ, AAn and MP

**Formal Analysis**: TW, AAm, PS, TK, JJ, AAn, and MP

**Funding Acquisition**: TW, AAm, and MP

**Investigation**: AAm, PS, TK, JJ, and MP

**Methodology**: AAn and MP

**Supervision**: MP

**Validation**: TW and MP

**Visualization**: TW and MP

**Writing – Original Draft Preparation**: TW and MP

**Writing – Review & Editing**: TW, AAn, and MP

## ACKNOWLEDGMENT

We thank Ryan Gentry Williams and Barbara Metzler for the English proof of this manuscript.

## SUPPORTING INFORMATION

**Table S1.** qRT-PCR primers used in this study.

**Table S2.** Bioinformatic analysis of AcuK and AcuM DNA binding motif (CCGN_7_CCG) in *T. marneffei* genome and their associated functions.

**Table S3.** List of genes involved in iron-sulfur cluster binding and oxidative phosphorylation, and their gene expression levels in the Δ*acuK* and Δ*acuM* vs wild type.

**Table S4.** List of selected genes involved in RIA, siderophore synthesis, gluconeogenesis and TCA cycle, and their gene expression levels in the Δ*acuK* and Δ*acu*M vs wild type.

**Table S5.** List of genes involved in ribosome, and ribosome biogenesis that are differentially reduced their expression levels in both mutants (log_2_fold-change ≤ −1 and p-value ≤ 0.05).

